# Model-guided Design of Biological Controller for Septic Wound Healing Regulation

**DOI:** 10.1101/2023.01.16.523937

**Authors:** Leopold N. Green, Pegah Naghshnejad, Derrick Dankwa, Xun Tang

## Abstract

Immune response is critical in septic wound healing. The aberrant and imbalanced signaling dynamics primarily cause a dysfunctional innate immune response, exacerbating pathogen invasion of injured tissue and further stalling the healing process. To design biological controllers that regulate the critical divergence of the immune response during septicemia, we need to understand the intricate differences in immune cell dynamics and coordinated molecular signals of healthy and sepsis injury. Here, we deployed an ordinary differential equation (ODE)-based model to capture the hyper and hypo-inflammatory phases of sepsis wound healing. Our results indicate that impaired macrophage polarization leads to a high abundance of monocytes, M1, and M2 macrophage phenotypes, resulting in immune paralysis. Using a model-based analysis framework, we designed a biological controller which successfully regulates macrophage dysregulation observed in septic wounds. Our model describes a systems biology approach to predict and explore critical parameters as potential therapeutic targets capable of transitioning septic wound inflammation toward a healthy, wound-healing state.

## Introduction

Wound healing is one of the most complex processes in multicellular organisms that involves a cascade of immune cells coordinated by molecular signals and aims to achieve tissue integrity, eradicate pathogen invasion, and promote tissue homeostasis. After an injury, platelets aggregate to form blood clots and prevent excessive blood loss in vascular tissues (Shannon 2020; Golebiewska and Poole 2015). The blood clots release various pro-inflammatory mediators, including tumor necrosis factor-alpha (TNF-α), to activate the bulk transmigration of neutrophils (*N*) into the tissue. Typically, neutrophils abundance steadily increases at the injury site for approximately 24 hours before undergoing apoptosis (Herrero-Cervera, Soehnlein, and Kenne 2022; M.-H. Kim et al. 2008). Neutrophil apoptosis releases cytokines and other molecules that recruit monocytes (*M0*), arriving within 5 to 6 hours post-injury (Marwick et al. 2018; El Kebir, Gjorstrup, and Filep 2012). The recruited monocytes differentiate into monocyte-activated macrophages. Macrophages are often considered the most important immune cell in the healing process, as they undergo phenotypic change towards the “classically activated” (*M1*) pro-inflammatory phenotype, followed by the “alternatively activated” (*M2*) anti-inflammatory phenotype. The macrophage plasticity (differentiation between *M0, M1*, and *M2*) is crucial in transitioning the wound microenvironment from a pro-inflammatory to a pro-resolution state (S. Y. Kim and Nair 2019; Krzyszczyk et al. 2018; Li et al. 2021). As inflammation is resolved, the *M2* anti-inflammatory macrophages secrete cytokine IL-4 and growth factors TGF-β, suppressing inflammatory immune cell activity while promoting extracellular matrix synthesis, recruiting fibroblasts, and stimulating mesenchymal cells to differentiate into myofibroblasts (Minutti et al. 2017).

In a healthy, acute wound-healing condition, inflammation is an essential step in the clearance of infection. Host immune white blood cells quickly enter the injury site to locate and remove pathogens. The immune response and transition from inflammation to proliferation and remodeling phases is typically tightly regulated, returning immune cell counts and their signal mediators to basal levels (Landén, Li, and Ståhle 2016). However, the dysregulated immune function produces a cytokine storm that impairs macrophage function, hindering the measures for preventing pathogen invasion—the consequences of a weakened immune response results in microenvironments ideal for systemic pathogenesis and sepsis.

Sepsis is a life-threatening condition initiated by pathogen infection and is one of the leading causes of morbidity and mortality worldwide (Rudd et al. 2020). Sepsis is fundamentally an immune-inflammatory condition characterized by two distinct phases in response to systemic infection: hyper-inflammation and prolonged immune suppression (Hotchkiss, Monneret, and Payen 2013; Nakamori, Park, and Shimaoka 2021). Excess production of inflammatory mediators such as TNF-α and IL-1β can further damage tissue and activate bacterial virulence. Concurrent immune suppression leads to macrophage paralysis, ineffectual wound healing, and increased susceptibility to infection.

Current efforts on inflammation regulation for chronic wounds and scar prevention have mainly focused on the delivery of growth factors, cytokines, and other immunomodulatory factors (Zhang and Ning 2021; Rudd et al. 2018; Tsirigotis et al. 2016). Despite early success, translating these therapies for septic wound resolution into the clinic has been difficult due to delivery methods and safety challenges (Ulloa et al. 2009; Vincent 2016; Malone and Schultz 2022). Most therapeutics currently rely on discrete doses, given via injections or applied topically to the wound site. Because of natural clearance, doses are given at supra-physiological levels to sustain drug presence throughout the wound healing process (Kumar and Singh 2015; Sjövall et al. 2018). This results in both safety concerns and poor long-term efficacy. Alternatively, dynamically regulating the immune system to suppress cytokine production and promote tissue repair and regeneration is an attractive approach for controlled and sustained drug release over longer periods, which is critical to the success of immune therapies.

In this work, we present a systems biology approach that deploys a model-based analysis framework to design a biological controller, to regulate bacterial invasion that often leads to local and systemic inflammation of septic wounds. Specifically, we developed a mechanistic model that captures the healing process based on literature findings and used local and global sensitivity analysis to identify the monitoring and regulation points along the pathway. We then proposed a biologically feasible controller (i.e., synthetic gene circuit) and demonstrated the efficacy of the controller by regulating various impaired healing conditions.

## Results

### Wound Healing Model Development

Recent systems medicine models reveal design principles of therapeutic circuit motifs that could improve impaired wound closure and excessive scar tissue (Cumming, McElwain, and Upton 2010; Flegg et al. 2015; Green et al. 2019; Adler et al. 2020). Computational models, including ordinary differential equation (ODE)-based models and agent based models, have demonstrated the capability of describing wound healing mechanisms while elucidating potential pathways and immune dynamics that may lead to therapeutic strategies (Walker et al. 2004; Mi et al. 2007; McDaniel et al. 2019; Vodovotz and An 2021; Zlobina, Xue, and Gomez 2022). In this work, we develop a simplified ODE model to describe the healing process in injured tissue, as outlined in Figure 2. The kinetic parameters are identified and tuned for the model to capture concurrent hyperinflammation and immune paralysis defined in septic wound healing.

**Figure 1.**
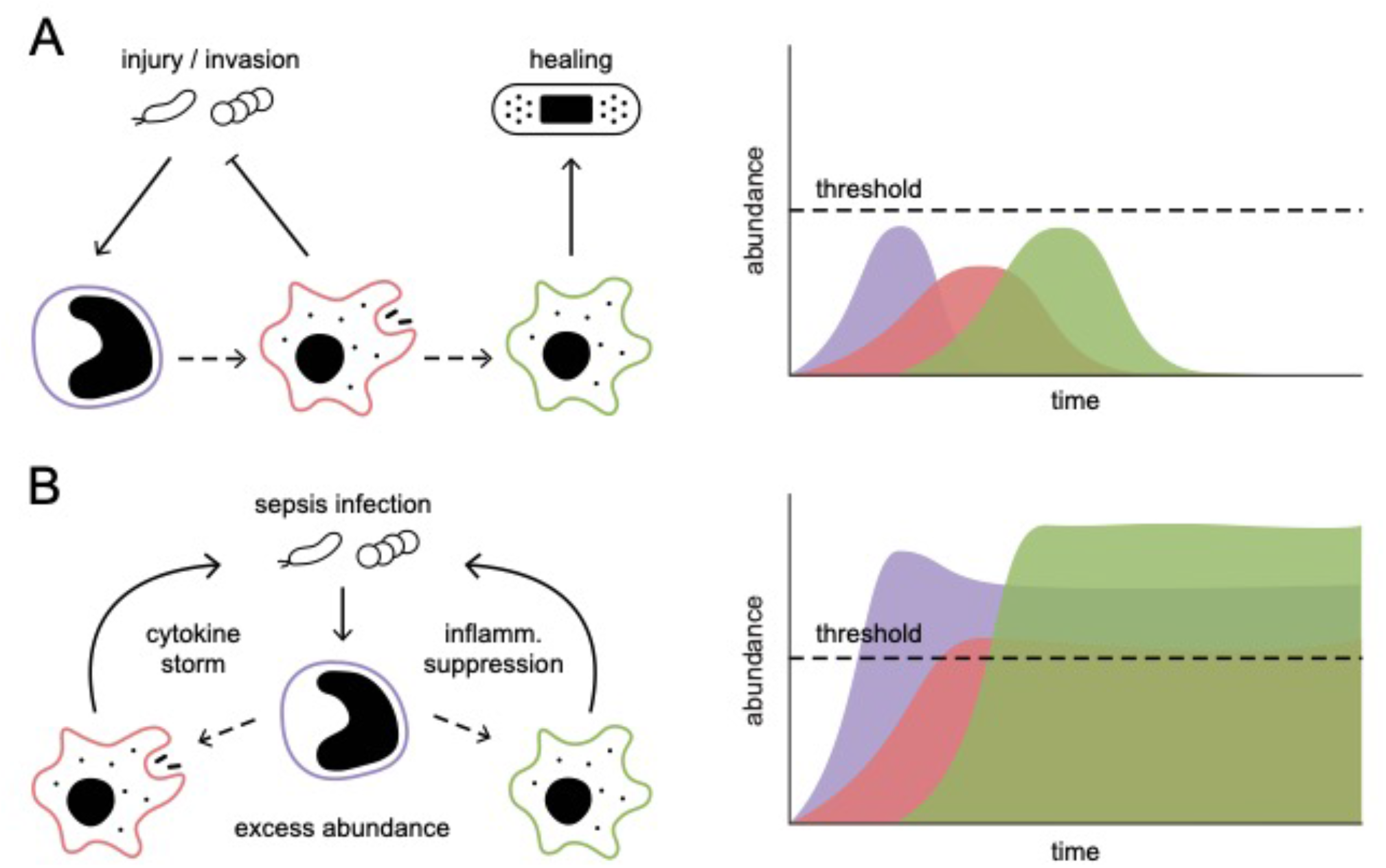
Diagram of macrophage polarization during wound healing. (A) balanced macrophage abundance and immune response eliminate disease and damaged tissue to promote homeostatic healing. (B) impaired macrophage polarization results in overabundance in all macrophage phenotypes. The cocureent cytokine storm and inflammation suppression lead to macrophage paralysis.

**Figure 2.**
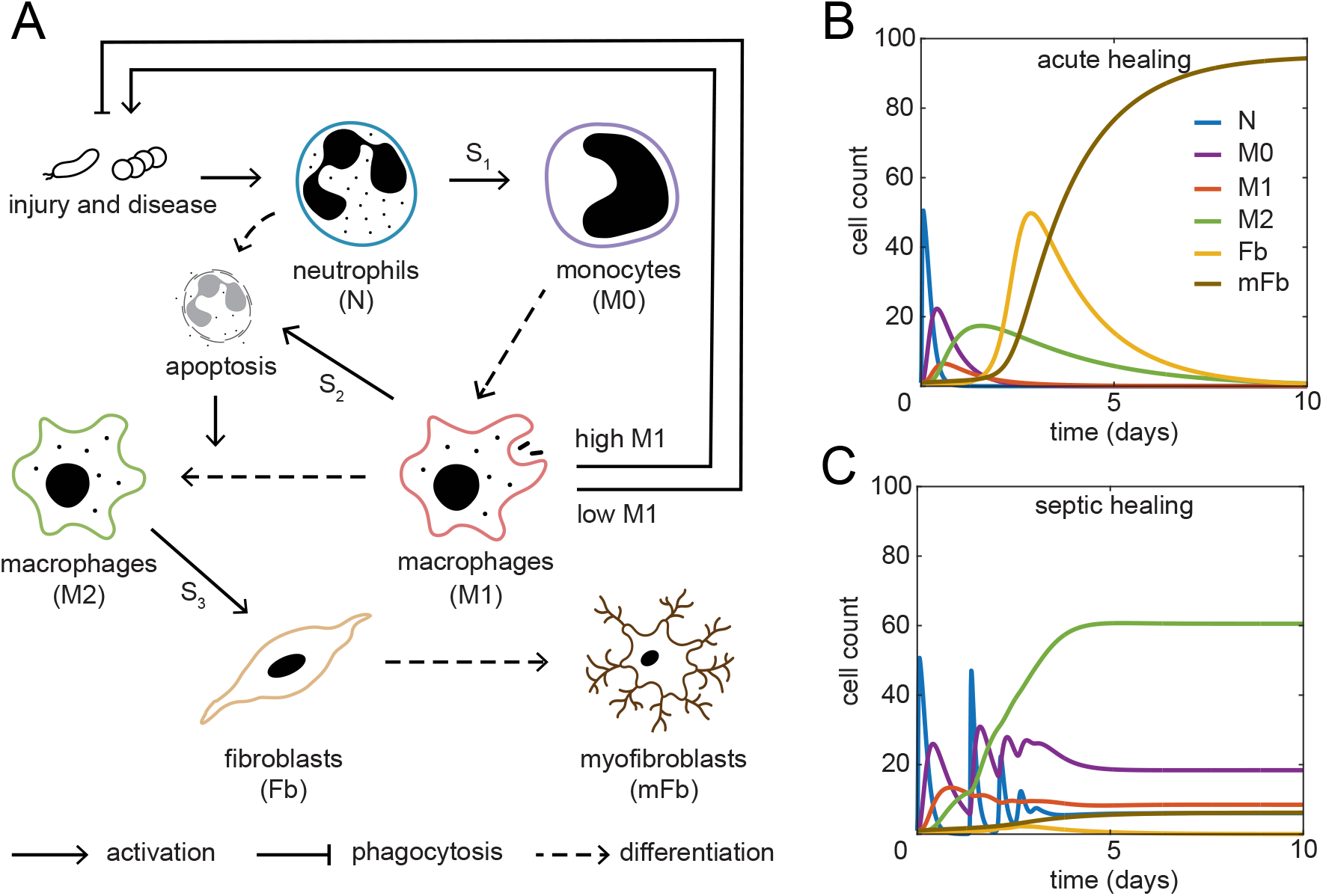
(A) Overview of wound healing immune cell dynamics following injury and pathogen invation. Simulation of (B) acute and (C) septic wound healing using parameters listed in Supplemental Table 1 and ODEs in equations (1-10).

Specifically, to facilitate the transition of each phase of wound healing based on cellular and molecular dynamics, we introduce chemical species S1, S2, and S3 to represent a class of critical cytokines and growth factors that coordinates healing. We also employ a Hill-type function to describe the impact of the injury on neutrophil recruitment. To represent the promoting and repressing effect of M1 on injury based on its relative abundance, we introduce a simple logic switch, *Effector*, defined as:

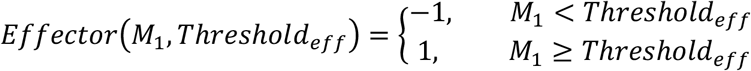

where, *M*_1_ is the cell count of the cellular species M1, and *Threshold*_*eff*_ is a tunable threshold to trigger the effector. When the M1 cell count is below the threshold, M1 will repress injury as expected during a healthy immune response; otherwise, M1 cells worsen injury. The detailed ODEs are given in equations 1-10. Where in general *α* represents the generation, recruiting rate of the corresponding species, *γ* represents induced apoptosis (γ_1_), differentiation (γ_2-5_), and cell recruiting (γ_6_) and rate, *μ* is the cell degradation rate, and *δ* is the signal degradation rate. *K* and *n* are the hill function constant. Capacity is defined as 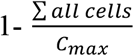, where *C*_*max*_ is the maximum number of cells allowed in the system, to introduce a constraint from the environment.

Using the derived ODE model, we first confirm if the model can capture the key characteristics of both the healthy and impaired wound healing processes. Figures 2B and 2C show the healthy and impaired cellular dynamics simulations during tissue injury, respectively. The kinetic parameter values are given in SI Table 1, and the simulations are solved with MATLAB ode23s solver. As indicated in Figure 2B, the healthy wound healing model demonstrates a sequential and timely cascade of cell dynamics of neutrophil, monocyte, M1, M2, fibroblast and ending with myofibroblast dominating the system after ten days. On the other hand, the cellular dynamics are distressed in the septic wound healing system (Figure 2C), with M0, M1, and M2 macrophage abundance remaining relatively high when compared to healthy healing dynamics. The observations verify that our ODE model can capture the fundamental features of healthy healing dynamics and impaired healing processes observed in sepsis disease.

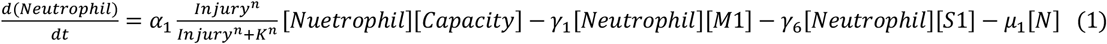

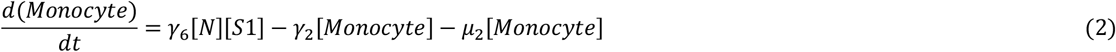

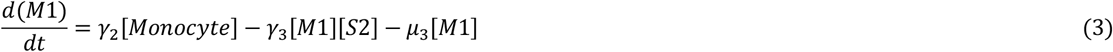

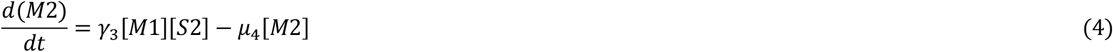

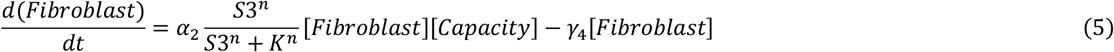

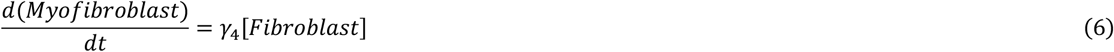

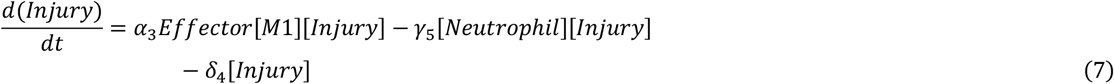

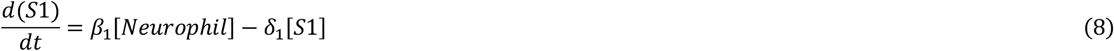

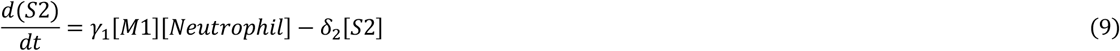

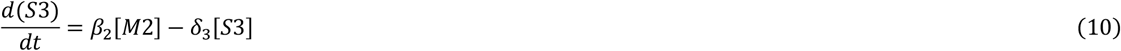

### Controller Design

After verifying the model, we introduced a controller that reverts the chronic healing dynamics toward a healthy, homeostatic immune response. Specifically, we designed a feedback controller capable of ensuring system stability, handling system uncertainty, and mitigating modeling errors to achieve a timely cascade of cellular dynamics ending with a high level of myofibroblast state after ten hours.

A typical feedback control involves 1) sensing, which measures the system’s status as an input to the controller, 2) control policy, a guideline to update the control input; and 3) actuation, which adjusts the manipulated variable according to the updated control input. Actively sensing and modulating immune cell and signal concentrations throughout the healing process is a viable therapeutic strategy, thus, we describe a proof-of-concept single-input-single-output controller to regulate vital immune processes. Our understanding of dysregulated immune response, altered signal concentrations, and impaired cellular function guided the controller design.

To identify the aforementioned components for our controller, we implemented a system’s framework outlined in Figure 3. Our model includes a local sensitivity analysis to identify the appropriate regulation point, i.e., manipulated variable to influence the process dynamics, and a global sensitivity analysis to identify the optimal cellular and molecular species to monitor and activate the feedback controller. Additionally, we use a biomolecular sequestration module as a threshold comparator to establish simple rules to trigger the controller when sepsis conditions arise.

**Figure 3.**
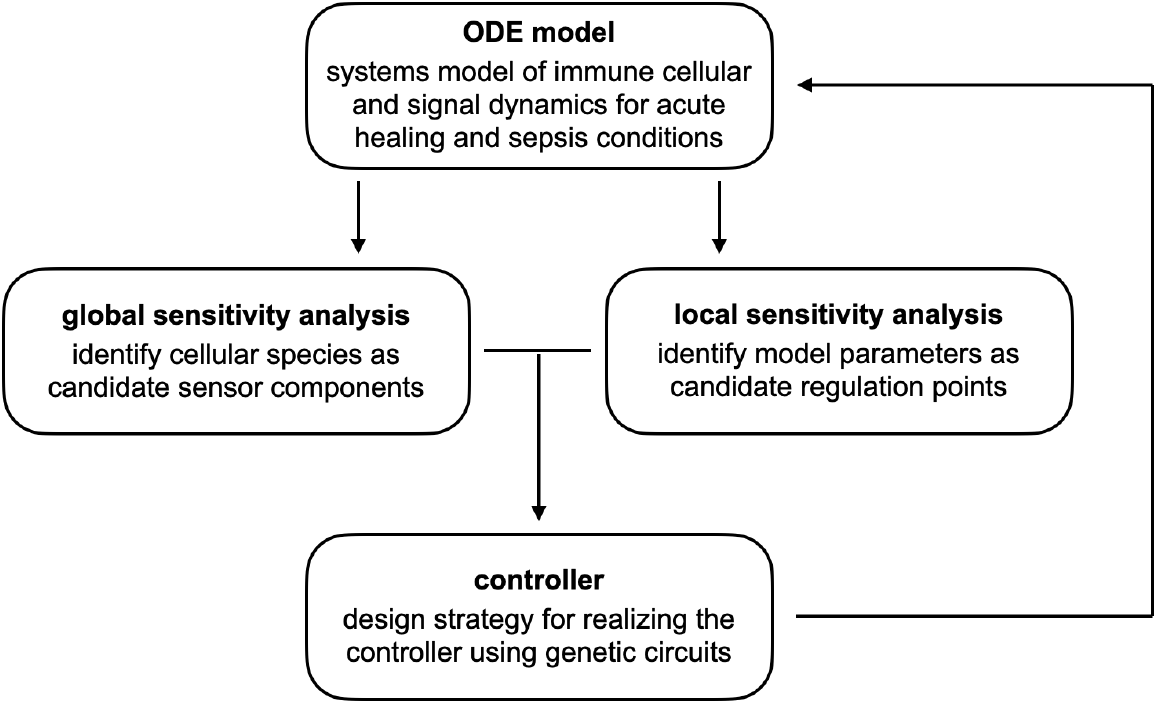
Framework for biological controller design and verification. Local sensitivity analysis provides information on the regulation point, and global snesitivity analysis identifies which cellular species to monitor, in the biological controller.

### Local Sensitivity Analysis for Regulation Chemical Identification

The local sensitivity analysis evaluates the model’s sensitivity to changes in a single parameter and provides insights into the most prominent variables that lead to impaired healing. Such parameters will serve as candidate regulation points for the controller to modulate.

In our study, we independently perturbed each of the 22 kinetic parameters to five discrete values spanning five orders of magnitude: [0.001, 0.01, 0.1, 1, 10] * *Nominal Value*. The parameter range represents a reasonable span of the biologically relevant values, while the coarse grain reduces the computational cost when covering such a broad range of values. Together we conducted a total of 22 × 5 = 110 simulations and analyzed the results using three metrics: Endpoint Cell Counts, Maximum Cell Counts, and Time to Peak. The “Endpoint Cell Count” quantifies the abundance of each cell species at the end of the 20-day simulation and determines if the system sequentially completes the phases of healthy healing or is halted in a chronic state. The “Maximum Cell Count” of each species and the “Time to Peak” metrics complement the “Endpoint Cell Count” by gauging the temporal signal and cellular dynamics to discern between healthy versus chronic healing processes.

Figure 4 summarizes the local sensitivity results based on the Endpoint Cell Count metric. The results for Maximum Concentration and Time to Peak are provided in SI Figure S1 and S2. For each parameter, we presented the average Endpoint Cell Count of the five simulations using the five values mentioned earlier for each of the six cell species. The error bars indicate the corresponding standard deviation. Results in Figure 4 suggest that the Endpoint Cell Count of fibroblast is least affected by the kinetic parameters. Conversely, the differentiation rate of fibroblast to myofibroblast, parameter *γ*_4_, shows a noticeable impact as illustrated by the large standard deviation. Myofibroblast displays the most sensitivity to single-parameter variation, as most parameters would affect the endpoint values quantified by the large standard deviation. In general, parameters *γ*_1_, *γ*_3_, *γ*_6_, *β*_1_, and *δ*_2_ show a significant impact on all the cell types. This observation further indicates that these parameters are candidate variables to regulate with the controller. Given the definitions of these parameters, we assigned *γ*_6_ as the manipulated variable, as it serves as a critical variable, upstream in the wound healing process: recruiting monocytes by way of neutrophils.

**Figure 4.**
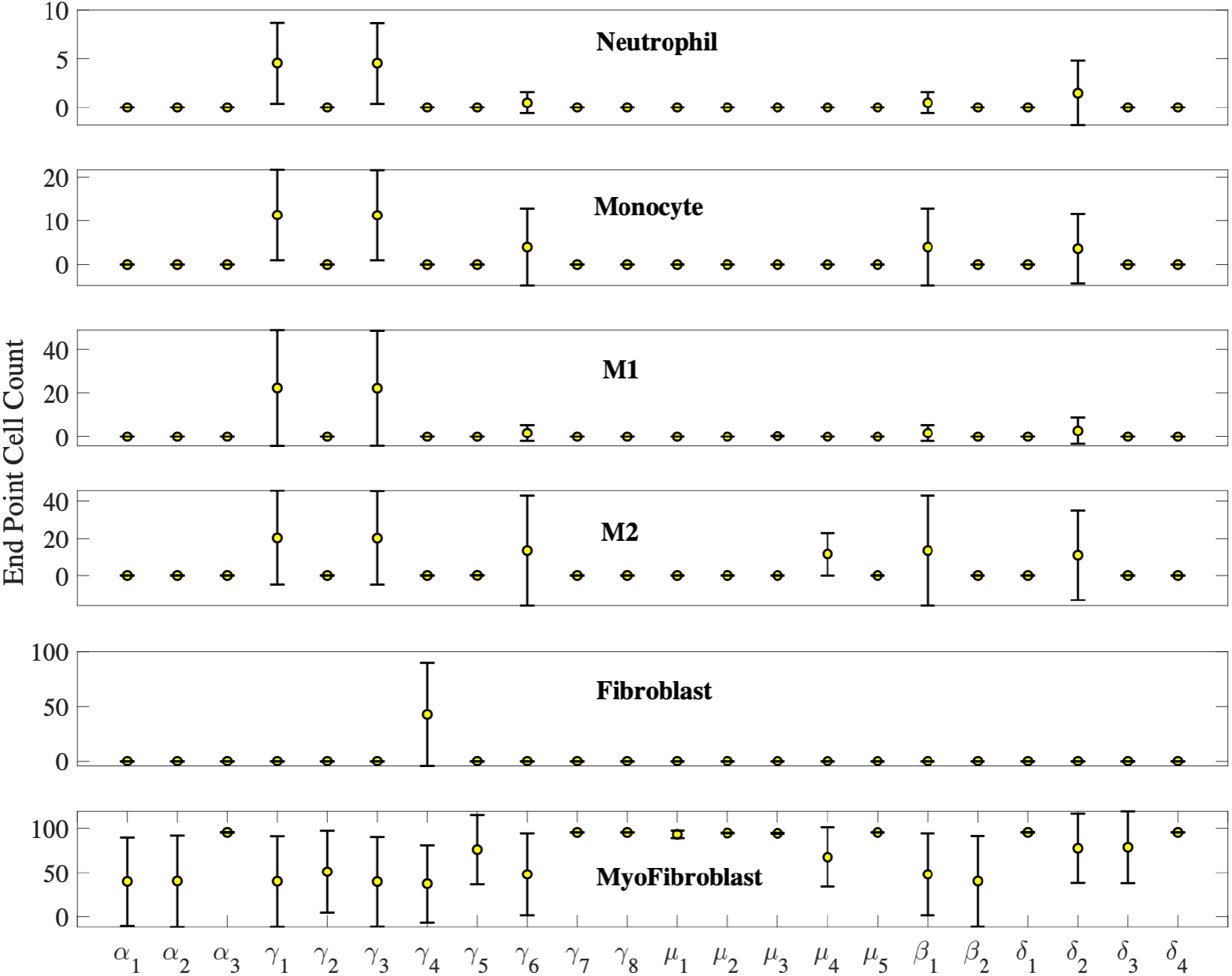
The endpoint cell count of the local sensitivity analysis reveals the dominating kinetic parameters in our system. Circles indicate the mean, and the error bar indicates the standard deviation of the endpoint concentration due to variations in each specific kinetic parameter.

### Global Sensitivity Analysis for Sensing Species Identification

While we identify the manipulated variable with local sensitivity analysis, we deploy global sensitivity analysis to identify the appropriate cell species for the controller to monitor. In the global sensitivity analysis, we randomly perturbed the most impactful kinetic parameters *γ*_1,_ *γ*_3,_ *γ*_6,_ *β*_1,_ and *δ*_2,_ within ±50% of their nominal values with a uniform distribution, and performed 1000 independent simulations to investigate the conditions that lead to chronic healing dynamics. This ±50% interval was chosen based on our observations from the local sensitivity analysis, which showed that 50% perturbations of the dominate parameters is sufficient in altering the healing dynamics toward chronic like-behavior.

Figure 5 summarizes the time trajectory of 1000 simulations of all the species, by which we group the simulations into two categories based on the Endpoint Cell Count: acute and chronic conditions. Specifically, when myofibroblast has the highest Endpoint Cell Count among all the cell types, with a count ≥ 40(selected as an example and is tunable based on the actual biological dynamics), the corresponding simulations are deemed as acute; otherwise, myofibroblast cell count below 40 cells is considered chronic healing. Such a criterion ensures that the most abundant cell type at the end of the process is myofibroblast, which corresponds to physiological healing conditions.

**Figure 5.**
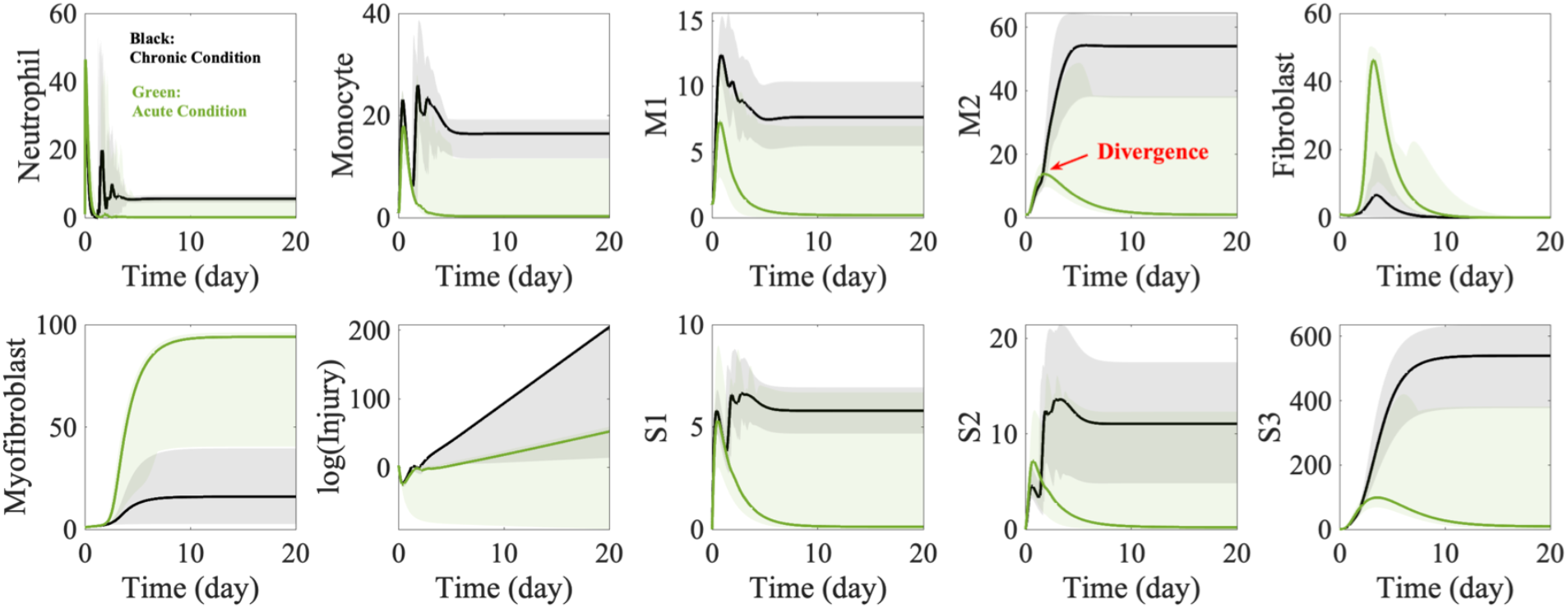
Global sensitivity analysis was performed to identify sensing points as feedback to the controller. Green lines represent the average acute condition simulation results, and black ones represent the averaged chronic condition simulation results, with the shaded areas indicating the entire space each condition has spanned in our simulation.

With this criterion, our modeled resulted in 43/1000 chronic and 957/1000 acute conditions. This observation further supports the selection of the nominal values with reasonable robustness to parameter uncertainties. The solid green and black lines in Figure 5 represent the average plots of the acute and chronic simulations, respectively, as the shaded region represents the entire range the dynamics covered for each simulation. Figure 5 shows that most chronic conditions were stalled due to high macrophage (M0, M1, M2) abundance, while the acute conditions demonstrated an ordered transition ending with maximum myofibroblast abundance. Comparing the difference between the acute and chronic healing conditions, we notice the time trajectory of the M2 cell count shows an evident divergence at around 15 cells at the early stage of the process. Therefore, we decide to monitor the M2 cell count as the feedback to trigger our controller.

### Threshold-based Single Input Single Output Biological Controller

The manipulated parameter in our controller, *γ*_6_, is the rate by which neutrophils recruit monocytes into the injured tissue. The controller’s objective is to confine the excessive accumulation of all macrophage phenotypes to avert the onset of chronic conditions. One control strategy is to limit the differentiation of M2 by down regulating the upstream events governed by the kinetic parameter *γ*_6_. Therefore, we propose the following simple logic regulator, capable of detecting the relative abundance of M2, *M*_*2*_, to a threshold, *Threshold*_*con*_:

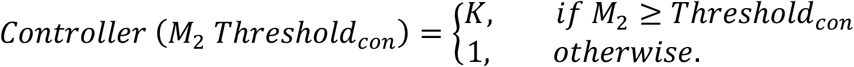

Where *K* is a large tunable number (“1), and we modify equation 2 as:

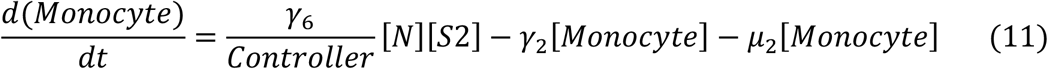

The design of the controller resets the total abundance of monocytes and macrophages prone to cause impaired healing conditions in the system by down-regulating neutrophil activation of monocytes. The controller design can be realized with a synthetic gene circuit, as shown in Figure 6. Specifically, we can use sequestration at the DNA, RNA, or protein level to establish a regulation threshold to activate the negative feedback in the system. The concentration of the negative inhibitor can shift the input-output threshold, while maintaining ultra-sensitivity, as shown in Figures 6 B and C (Buchler and Cross 2009).

**Figure 6.**
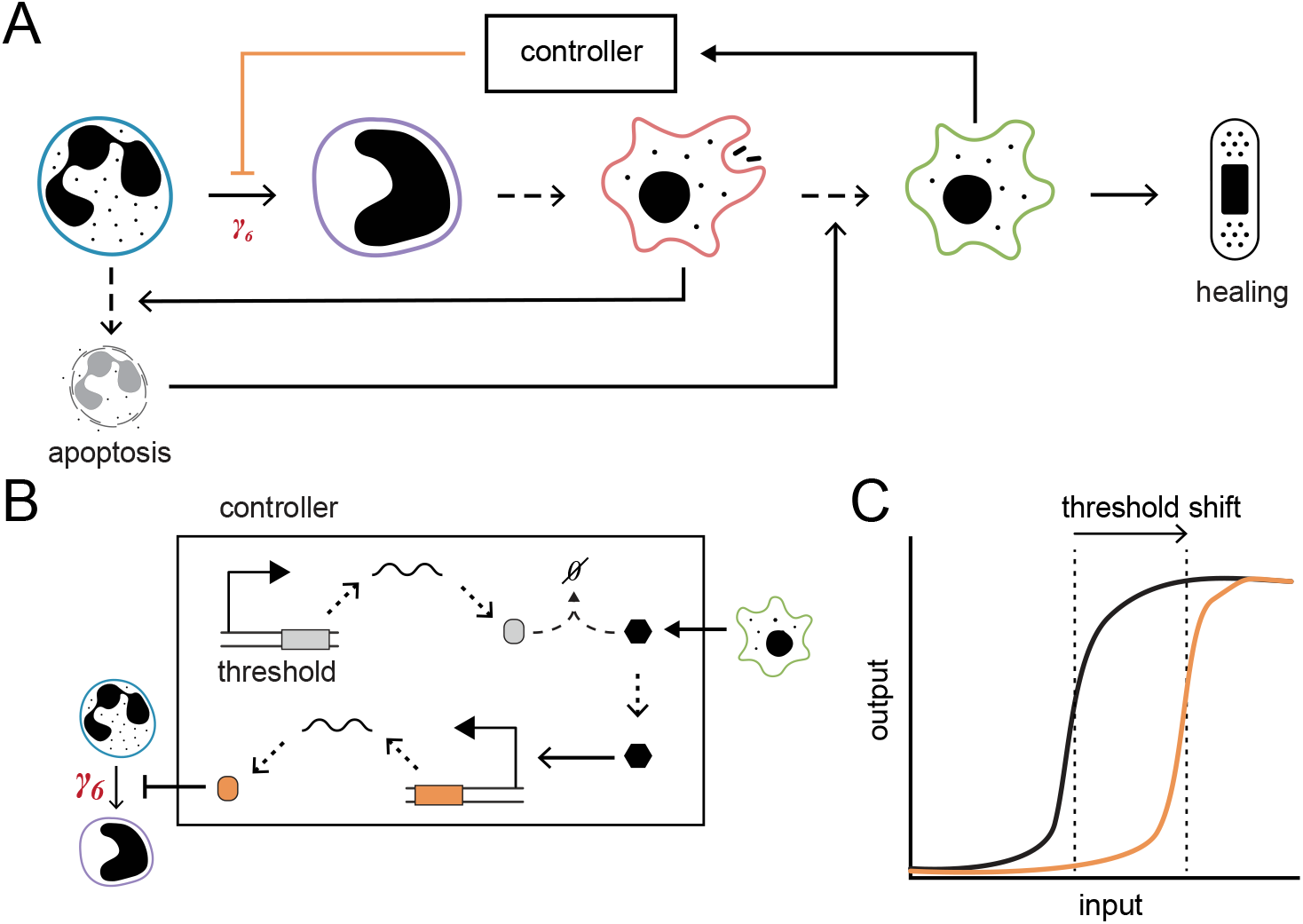
(A) Schematic of the controller integrated into the cellular imune circuit. (B) Realization of controller design using sequestration gene circuit to establish a threshold setpoint. The controller output (orange) is activated when the M2 input signal (black) is above the threshold comparator compound (gray). (C) Illustration of input-output dynamics with increased threshold setpoint.

Two factors would affect the controller performance: the concentration of M2 to trigger threshold, *Threshold*_*con*_ and the parameter *K* value. From our sensitivity analysis, varying these two parameters (SI Figure S3) indicates that the performance of the controller is dominated by the threshold value and is less sensitive to the value of *K*. This observation is expected, as the threshold determines the controller’s responsiveness to the system dynamics. For illustration purposes, we present results for *K* = 100, an *Threshold*_*con*_ = 20, for analysis.

### Controller Performance Evaluation

To evaluate the performance of our controller, we implemented the controller on all the simulated trajectories from the global sensitivity analysis in Figure 6. We anticipate that the controller will revert the chronic healing dynamics and deliver an improved healing process while not interfering with a healthy healing process.

Results in Figure 7 summarize the controlled (red) and uncontrolled (black) results of the 43 chronic conditions obtained from the global sensitivity analysis. The solid black and red lines indicate the average dynamics of the uncontrolled and the controlled simulations, respectively. The shaded area represents the range covered space by the simulations. Figure 7 illustrates that the controller reverts all chronic conditions to healthy by down regulating the cell count of monocyte, M1, and M2, which subsequentially lead to the max (or near max) abundance of myofibroblast at the end of the simulation. One noticeable point is that the controller introduced oscillations into the system, exemplified by neutrophils, monocyte, S1, and S2, before improving the dynamics of each cell type. A comparison of the log scale injury profile demonstrates that injury explodes over time without control, as observed in septic wounds, but with the controller, injury is reduced allowing for healthy healing. An example of individual controlled and uncontrolled simulations is provided in SI Figure S4, which shows similar observations as the statistically averaged dynamics.

**Figure 7:**
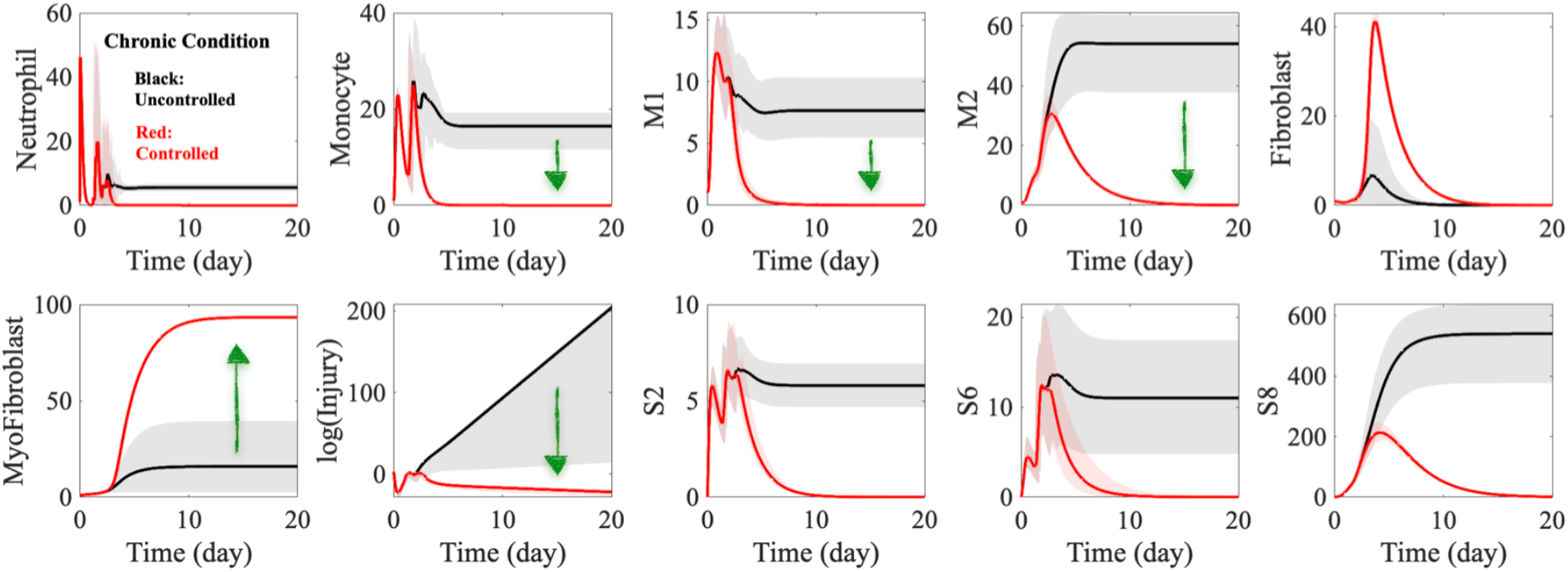
Comparison of controlled and uncontrolled simulations. Statistical results of the controlled and uncontrolled chronic conditions for each cell and signal. The black and red lines represent the uncontrolled and uncontrolled average dynamics. Simulation results suggest the controller can revert the chronic healing condition to acute wound healing. B) Statistical results of the controlled and uncontrolled acute conditions for each cell and signal. The green and red lines represent the uncontrolled and uncontrolled average dynamics. Simulation results suggest no significant interference from the controller to the healthy healing process.

Out of the 957 acute simulations from the global sensitivity analysis, 97% of the simulations ended up with an endpoint myofibroblast cell count of ≥ 90, with the remaining 3% of simulations with an endpoint myofibroblast cell count of ≤ 60, as demonstrated in the histogram distribution in Figure 8A. To evaluate the impact of the controller on the acute healing processes, we applied the controller to two representative simulations, one with a high endpoint myofibroblast and one with a low myofibroblast, as shown in Figure 8B. The comparison reveals that the controller does not interfere with the system dynamics during the acute healing process (Scenario 1), while the controller improves impaired healing during sepsis conditions (Scenario 2). Indeed, the Scenario 2 condition could be classified as sepsis in wound healing.

**Figure 8.**
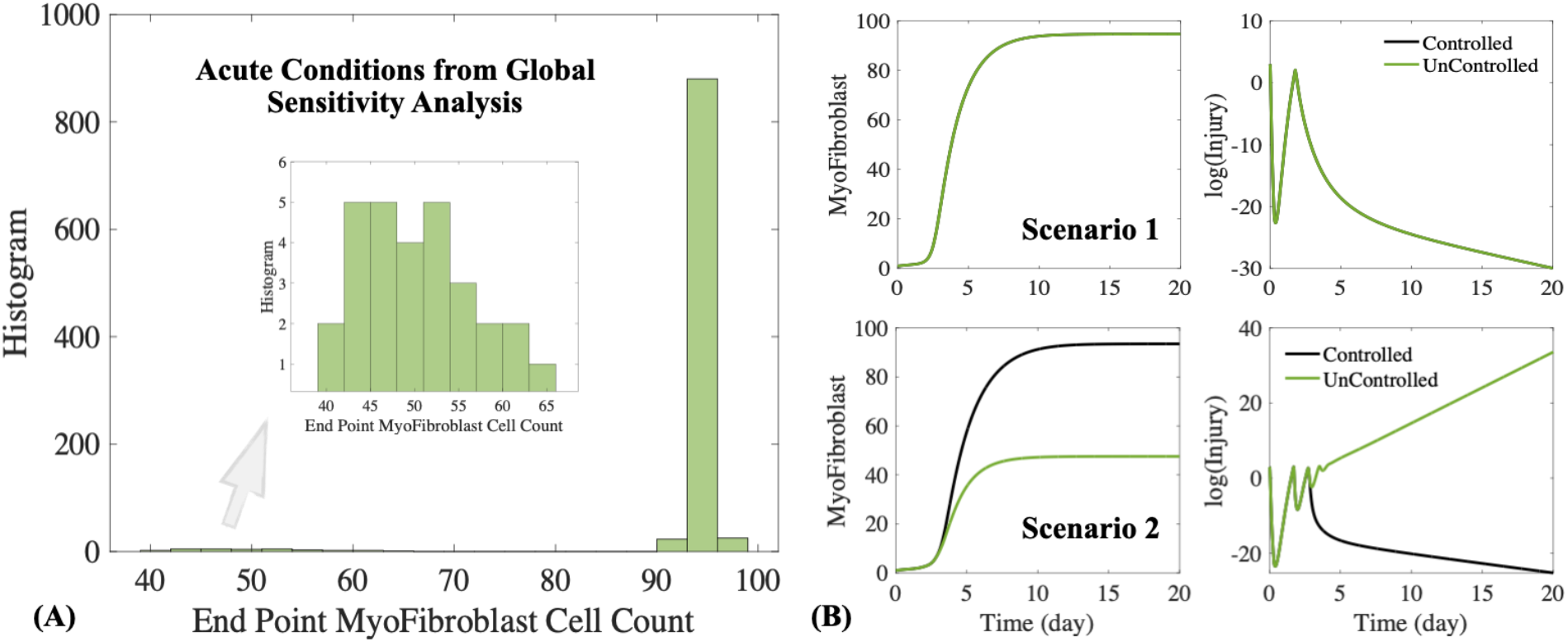
Evaluation of controller interferences on acute healing conditions. (A) histogram of endpoint myofibroblast cell count of the 957 acute condition simulations from the global sensitivity analysis. (B) Two different acute healing scenarios suggest the controller impact on the system depends on the “wellness” of the system, where no impact is observed if the system is in a good acute healing condition (Scenario 1). The green plots are with the controller, and the black plots are without the controller.

Altogether, the simulation results confirm that our controller can improve the chronic healing condition and has minimum impact on the system if it is in an acute healing condition.

## Discussion

Bacterial infections leading to sepsis are a key challenge facing our global healthcare system. The pathophysiology of sepsis is vaguely defined as the dysregulated host response to infection. Sepsis is traditionally considered a biphasic condition where immunosuppression follows hyperinflammation. However, recent studies describe the underlying mechanisms to include concurrent hyperinflammation and immunosuppression from the onset of sepsis. Sepsis treatment depends mainly on candid measures, including source control, resuscitation, and palliative interventions (Mackenzie and Lever 2007; Dugar, Choudhary, and Duggal 2020). There remains a growing need for early diagnosis and responsive treatments at the onset of septicemia and impaired wound healing caused by the septic invasion. An ideal septic wound therapeutic must maintain a balanced control of the hyper and the hypo-inflammatory phase in a dysfunctional immune process and prevent septicemia and further organ malfunction (Tsirigotis et al. 2016). Systems biology models are valuable tools to elucidate critical signaling pathways and design potential controller motifs that may serve as immune-modulating interventions.

In this work, we developed a set of ordinary differential equations that captures the prominent characteristics of sepsis’s impaired immune cellular dynamics. Our systems biology model also illustrates how dysregulated macrophage count, particularly the concurrent overabundance of M0, M1 and M2 macrophages, results in poor wound healing, leading to recurrent wound infections. While the model explains the severe derangement of immune cell dynamics, it also highlights an ideal monitoring and regulation point to reset immune paralysis based on local and global sensitivity analysis. Furthermore, the proposed feedback biological controller can recover the septic inflammatory condition to the healed state by modulating the neutrophil to M0 differentiation rate in response to M2 dynamics.

Cytokines and growth factors are critical mediators as they are involved in crucial transition points during the immune process. For example, TGF-β is responsible for resolving inflammation, skewing the polarization of M2 macrophage phenotype, recruiting fibroblasts, and stimulating mesenchymal cells to differentiate into myofibroblasts. Our local and global sensitivity analysis illustrated that pro and anti-inflammation signaling pathways remained upregulated during pathologic localization and infusion observed in sepsis inflammation (Nedeva, Menassa, and Puthalakath 2019; Qiu, Liu, and Zhang 2019). The controlled results suggest that detecting excess cell and signal abundance concerning a critical threshold, thus activating the regulation of the key transition points, is a viable and practical approach to dynamically controlling the healing process.

Computational models are generally limited by assumptions that simplify the complexity of biological systems. The work presented here highlights a primary controller for subverting sepsis wound healing based on the offset of immune dysregulation and suppression. Our model focuses only on the impaired immune responses of the predominant innate cells in wound healing: neutrophil, macrophage, fibroblast, and myofibroblast, and overlooks deficits in the adaptive immune response, which are also critical in septic-induced mortality. Like the innate immune response, the adaptive cell types, including T cells, B cells, and dendritic cells, play immunoregulatory roles in mitigating tissue damage and facilitating immune homeostasis after infection or injury (Brady, Horie, and Laffey 2020). We expect such simplification to render discrepancy in experimental implementation and anticipate further modification of the model to reflect experimental details for control performance.

Additionally, we used single molecular elements to represent a class of immune cytokine and chemokine signals to simplify the complex yet essential pleiotropic nature of immune signals in our model. In our model and analysis, we also used cellular and signal parameter values to generalize impaired wound healing during sepsis. In future work, we will identify the critical biomarkers responsible for pivotal transition points defined in our model (e.g., neutrophil apoptosis, macrophage polarization) and characterize the parameter values for signal production and secretion rates from the predominant cell type. As more information becomes available, we will modify our model assumptions and incorporate detailed mechanisms, to improve the relevancy and accuracy of the model and the inferred pathogenesis of sepsis.

The framework presented in this computational study provides a potentially viable approach to designing a biological regulator for a complex biological pathway. We expect our work to guide experimental exploration to advance our understanding of the immune dynamics in sepsis and to facilitate the development of synthetic-based theragnostic for septic wound healing.

## Materials and Methods

The mechanistic ODE model was developed around the key species in the system, following the low of mass action. All the simulations presented in this manuscript were conducted in MATLAB_R2022a, and solved with ode23s solver. For the local sensitivity analysis, we independently perturbed each of the 22 kinetic parameters in our model to five discrete values, spanning five orders of magnitude: [0.001, 0.01, 0.1, 1, 10] * *Nominal Value*. This parameter range represents a reasonable span of the biologically relevant values, while the coarse grained interval reduces the computational cost when covering such a broad range of values. The nominal value for each parameter is provided in SI Table 1. Together, we conducted a total of 22 × 5 = 110 simulations and analyzed the results using the three metrics: Endpoint Cell Counts, Maximum Cell Counts, and Time to Peak, as defined in the manuscript. In the global sensitivity analysis, we randomly perturbed the most impactful kinetic parameters *γ*_1,_ *γ*_3,_ *γ*_6,_ *β*_1,_ and *δ*_2,_ within ±50% of their nominal values with a uniform distribution with MATLAB command *rand*, and performed 1000 independent simulations to investigate the conditions that lead to chronic healing dynamics. This ±50% interval was chosen based on our observations from the local sensitivity analysis, that 50% perturbations of the dominate parameters is sufficient in altering the healing dynamics toward chronic like-behavior. To evaluate the controller performance, 1000 simulations with the controller were conducted using the same kinetic parameters used for the global sensitivity analysis. All the results plots were generated in MATLAB. Codes and scripts are available per reasonable request.

## Author Contribution

L. G. and X. T. conceptualized the idea, developed the model, and designed the work. P. N. and X. T. performed the simulation. L. G., P. N. and X. T. analyzed the data. All authors contributed to the writing.

## Acknowledgments

L. G. and D. D. are supported by ONR grant #N00014-21-1-2624 and by the startup fund provided by the Weldon School of Biomedical Engineering at Purdue University. X. T. and P. N. are supported by NSF grant #2223720 and the startup fund provided by the Cain Department of Chemical Engineering at Louisiana State University.

