## Supplementary Information for "Model-guided Design of Biological Controller for Septic Wound Healing Regulation"

Leopold N. Green[*,1], Pegah Naghshnejad[2], Derrick Dankwa [1], Xun Tang[*,2]

[1] Weldon School of Biomedical Engineering, Purdue University, West Lafayette, IN 47905
[2] Cain Department of Chemical Engineering, Louisiana State University, Baton Rouge, LA 70803
[*] Corresponding authors

**Supplementary Table S1.** Nominal Kinetic Parameters for our Simulations. The parameters are varied between 10*Nominal values and 0.001*Nominal values, for local sensitivity analysis to account for the effects of changing each value.

| Parameter | Nominal value | Lower Bound | Upper Bound | Unit | Description |
|---|---|---|---|---|---|
| n | 1 | 1 | 1 | - | Hill function coefficient |
| k | 200 | 200 | 200 | - | Hill function coefficient |
| $Threshold_{eff}$ | 0.05 | 0.05 | 0.05 | M | Threshold for Injury regulation by M1 |
| $C_{max}$ | 100 | 100 | 100 | M | Maximum number of cells |
| $\alpha_1$ | 2000 | 2 | 20000 | $Sec^{-1}$ | production of neutrophil |
| $\alpha_2$ | 20 | 0.02 | 200 | $Sec^{-1}$ | production of Fibroblast |
| $\alpha_3$ | 2.5 | 0.0025 | 25 | $Sec^{-1}$ | production of Injury |
| $\gamma_1$ | 0.25<br>0.288 (Figure 2A)<br>0.167 (Figure 2B) | 0.00025 | 2.5 | $Sec^{-1}$ | induced apoptosis of Neutrophil via M1 |
| $\gamma_2$ | 1.5 | 0.0015 | 15 | $Sec^{-1}$ | differentiation rate from Monocyte to M1 |
| $\gamma_3$ | 0.375<br>0.510 (Figure 2A)<br>0.275 (Figure 2B) | 0.000375 | 3.75 | $Sec^{-1}$ | differentiation rate from M1 to M2 |
| $\gamma_4$ | 0.75 | 0.00075 | 7.5 | $Sec^{-1}$ | differentiation rate from Fibroblast to Myofibroblast |
| $\gamma_5$ | 2.5 | 0.0025 | 25 | $Sec^{-1}$ | induced apoptosis of Injury via Neutrophil |
| $\gamma_6$ | 0.75<br>0.810 (Figure 2A)<br>0.953 (Figure 2B) | 0.00075 | 7.5 | $Sec^{-1}$ | differentiation of Neutrophil to M0 |
| $\gamma_7$ | 2.5 | 0.0025 | 25 | $Sec^{-1}$ | induced apoptosis of Injury via M1 |
| $\gamma_8$ | 2.5 | 0.0025 | 25 | $Sec^{-1}$ | induced apoptosis of N via M1 |
| $\mu_1,$ | 0.4 | 0.0004 | 4 | $Sec^{-1}$ | Neutrophil removal cell |
| $\mu_2$ | 0.4 | 0.0004 | 4 | $Sec^{-1}$ | Monocyte removal cell |
| $\mu_{3,}$ | 0.4 | 0.0004 | 4 | $Sec^{-1}$ | M1 removal cell |
| $\mu_4$ | 0.4 | 0.0004 | 4 | $Sec^{-1}$ | M2 removal cell |
| $\mu_5$ | 0.4 | 0.0004 | 4 | $Sec^{-1}$ | Fibroblast removal cell |
| $\beta_1$ | 0.5<br>0.307 (Figure 2A)<br>0.589 (Figure 2B) | 0.0005 | 5 | $Sec^{-1}$ | Production of S2 |
| $\beta_2$ | 6 | 0.006 | 60 | $Sec^{-1}$ | Production of S8 |
| $\delta_1$ | 0.6 | 0.0006 | 6 | $Sec^{-1}$ | S2 degradation |
| $\delta_2$ | 0.6<br>0.830 (Figure 2A)<br>0.830 (Figure 2B) | 0.0006 | 6 | $Sec^{-1}$ | S6 degradation |
| $\delta_3$ | 0.6 | 0.0006 | 6 | $Sec^{-1}$ | S8 degradation |
| $\delta_8$ | 0.5 | 0.0005 | 5 | $Sec^{-1}$ | Injury degradation |
| K | 100 | - | - | - | K value for controller |
| $Threshold_{con}$ | 20 | - | - | M | Controller threshold |

**Supplementary Figure S1.** Local sensitivity analysis in terms of maximum cell count. The maximum cell count from the local sensitivity analysis for each parameter is analyzed to show the impact of single parameter variation on the maximum amount of cells can be reached during the simulation. Circles indicate the mean, and the error bar indicates the standard deviation of the maximum cell count for each parameter. Results indicate neutrophil and M1 show relatively low sensitivity to single parameter perturbation, while most of the kinetic parameters showing noticeable impact on all the other cell types. Specifically, parameters $\alpha_1, \gamma_1, \gamma_2, \gamma_3, \gamma_5, \gamma_6, and\ \beta_1$ show the most significant impact on most of the cell types.

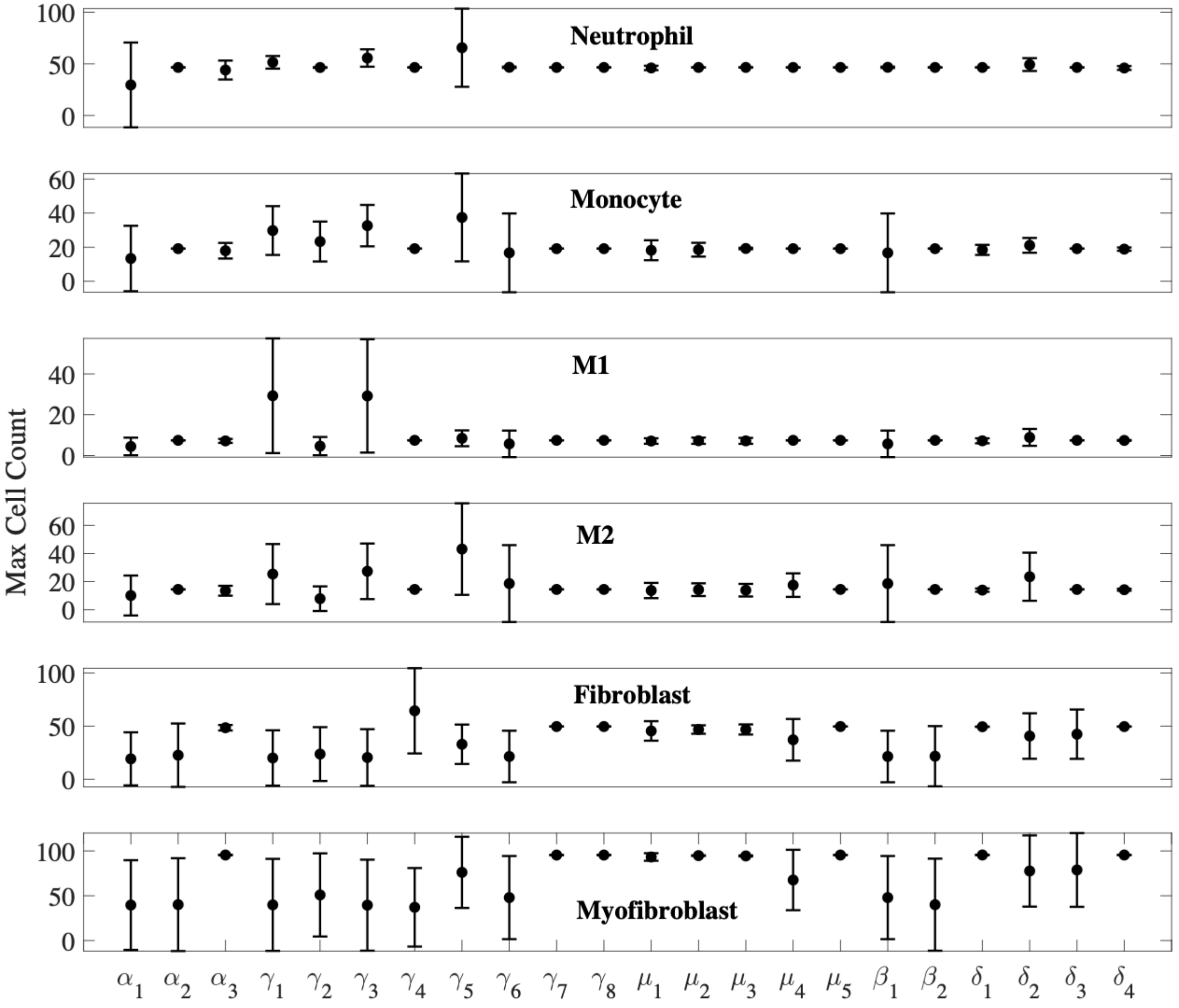

**Supplementary Figures S2.** Local sensitivity analysis in terms of time to peak. The take taken for each cell type to reach its maximum cell count in each simulation, from the local sensitivity analysis for each parameter, is analyzed to show the impact of single parameter variation on the temporal dynamics of the process. Circles indicate the mean, and the error bar indicates the standard deviation of the maximum cell count for each parameter. Results indicate parameters $\gamma_1, \gamma_3, \delta_2$ show the most significant impact on most of the cell types, while $\gamma_6, \beta_1$ also demonstrate noticeable impact to the system.

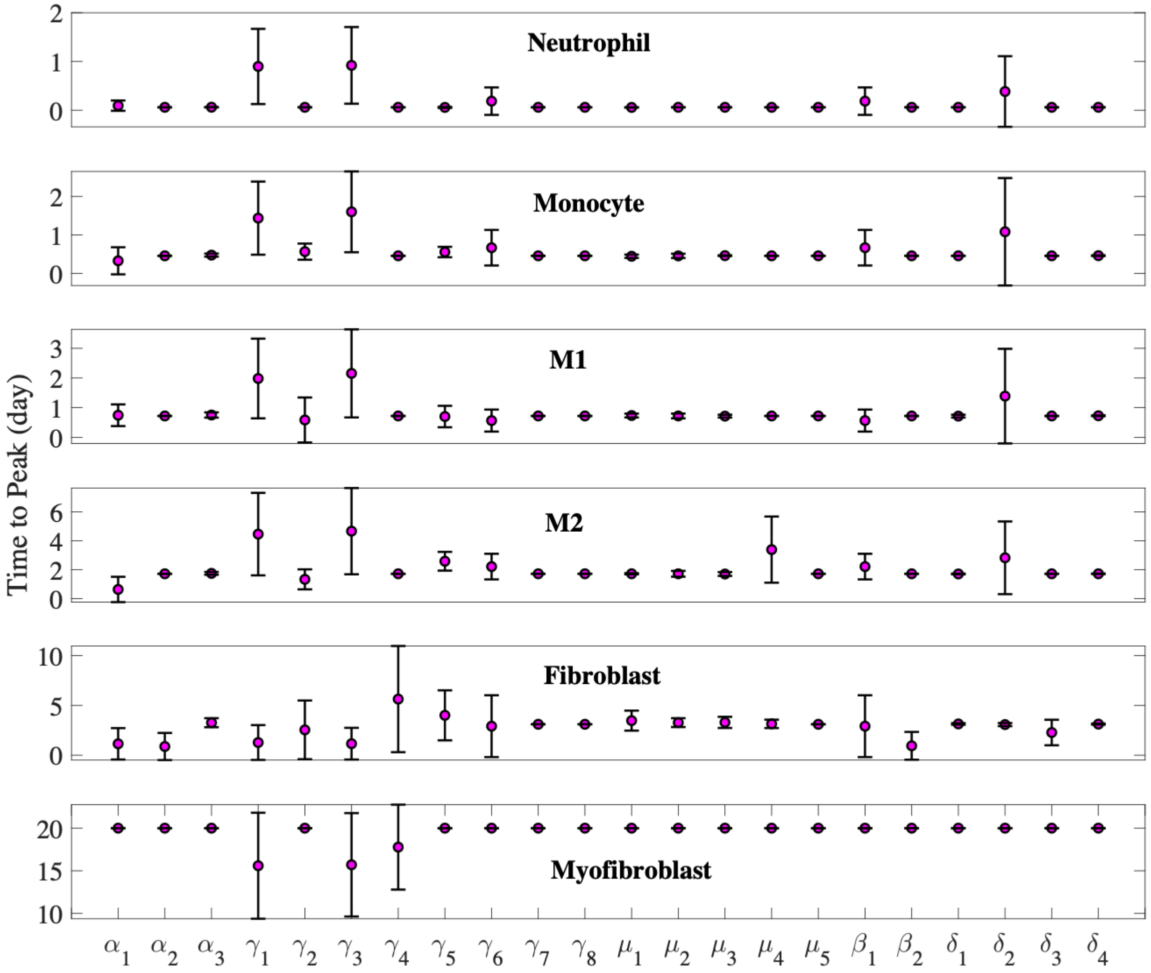

**Supplementary Figures S3.** Controller sensitivity analysis with respect to *K* value and $Threshold_{con}$, on controlling the unhealthy simulations from the global sensitivity analysis. The results reveal that the threshold for M2 cell count used to trigger the controller, dominates the dynamics rather than the *K* value, which is how much of the parameter $\gamma_6$ is to be regulated.

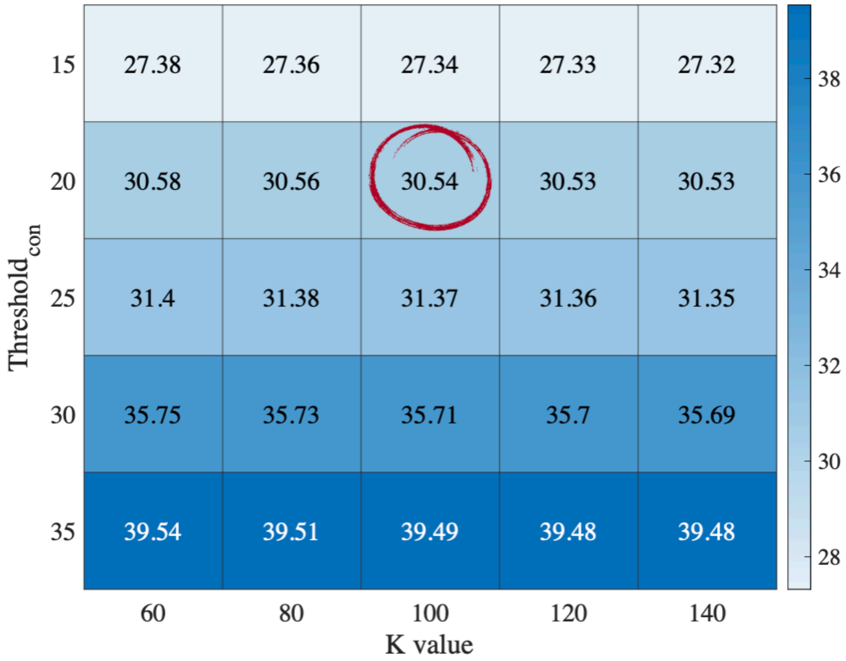

**Supplementary Figures S4.** Comparison of a single controlled and uncontrolled chronic condition simulation, indicating similar observations from the statistical analysis in Figure 7. The controller is able to direct the system dynamics from chronic condition to acute healing process, with oscillation dynamics introduced to the system, especially in neutrophil, monocyte, M1, injury, and chemicals S1, and S2.

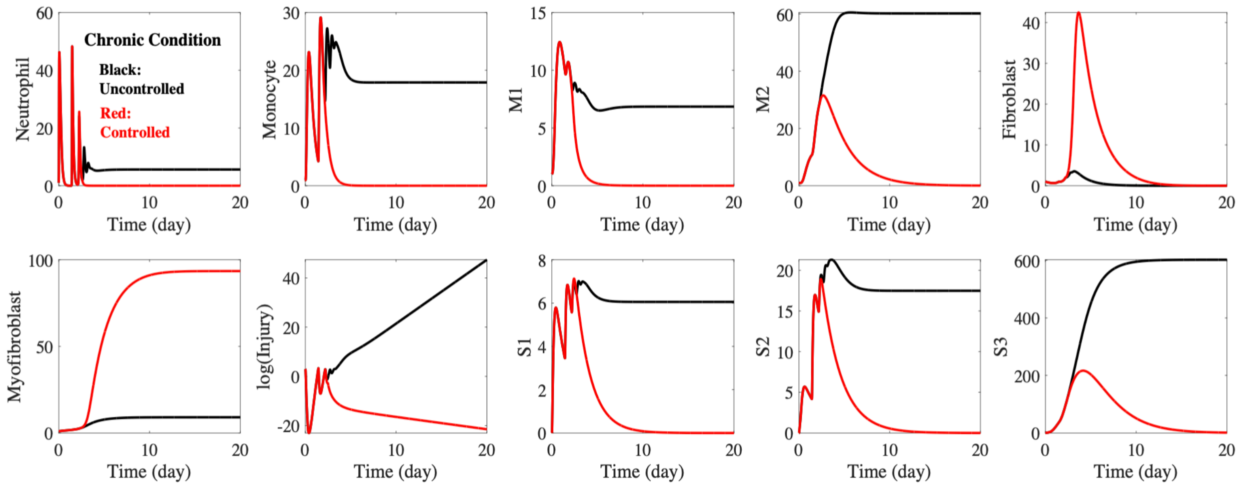
